# Metabolic reconstitution by a gnotobiotic microbiota varies over the circadian cycle

**DOI:** 10.1101/2021.09.08.456534

**Authors:** Daniel Hoces, Jiayi Lan, Sun Wenfei, Tobias Geiser, Elisa Cappio Barazzone, Markus Arnoldini, Sven Nowok, Andrew J Macpherson, Bärbel Stecher, Renato Zenobi, Wolf-Dietrich Hardt, Christian Wolfrum, Emma Slack

**Affiliations:** Laboratory for Food Immunology, Institute of Food, Nutrition and Health, Department of Health Sciences and Technology, ETH Zürich, 8093 Zürich, Switzerland; Laboratory of Organic Chemistry, Department of Chemistry and Applied Biosciences, ETH Zürich, 8093 Zürich, Switzerland; Laboratory of Translational Nutrition Biology, Institute of Food, Nutrition and Health, Department of Health Sciences and Technology, ETH Zürich, 8603 Schwerzenbach, Switzerland; ETH Phenomics Center, Department of Biology, ETH Zürich, 8093 Zürich, Switzerland; Department of Biomedical Research, Gastroenterology & Mucosal Immunology, University of Bern, 3010 Bern, Switzerland; Max-von-Pettenkofer Institute, LMU Munich, Munich, Germany; German Center for Infection Research (DZIF), partner site LMU Munich, 80539 Munich, Germany; Institute of Microbiology, Department of Biology, ETH Zürich, 8093 Zürich, Switzerland

**Author notes:** Stanford University, Department of Bioengineering, CA 94305, USA.

**Keywords:** Microbiota, Host Metabolism, Gnotobiotic, Circadian rhythm.

## Abstract

The capacity of the intestinal microbiota to degrade otherwise indigestible diet components is known to greatly improve the recovery of energy from food. This has led to the hypothesis that increased digestive efficiency may underlie the contribution of the microbiome to obesity. OligoMM12-colonized gnotobiotic mice have a consistently higher fat-mass than germ-free or fully colonized counterparts. We therefore investigated their food intake, digestion efficiency, energy expenditure and respiratory quotient using a novel isolator-housed metabolic cage system which allows long-term measurements without contamination risk. This demonstrated that microbiota- released calories are perfectly balanced by decreased food intake in fully colonized versus gnotobiotic OligoMM12 and germ-free mice fed a standard chow diet, i.e., microbiota-released calories can in fact be well-integrated into appetite control. We also observed no significant difference in energy expenditure per gram lean mass between the different microbiota groups, suggesting that cumulative very small differences in energy balance, or altered energy storage must underlie fat accumulation in OligoMM12 mice. Consistent with altered energy storage, major differences were observed in the type of respiratory substrates used in metabolism over the circadian cycle: in germ-free mice the respiratory exchange ratio was consistently lower than that of fully colonized mice at all times of day, indicative of more reliance on fat and less on glucose metabolism. Intriguingly the RER of OligoMM12-colonized gnotobiotic mice phenocopied fully colonized mice during the dark (active/eating) phase but phenocopied germ-free mice during the light (fasting/resting) phase. Further, OligoMM12-colonized mice showed a germ-free-like drop in liver glycogen storage during the light cycle and both liver and plasma metabolomes of OligoMM12 mice clustered closely with germ-free mice. This implies the existence of microbiota functions that are required to maintain normal host metabolism during the resting/fasting phase of circadian cycle, and which are absent in the OligoMM12 consortium.

## INTRODUCTION

The gut microbiota is currently considered a key regulator of host energy metabolism (Sonnenburg and Bäckhed, 2016). In the absence of a microbiota, mice accumulated less fat (Bäckhed et al., 2004) and were protected from obesity induced by certain types of high-fat diets (Bäckhed et al., 2007; Fleissner et al., 2010; Kübeck et al., 2016). Several mechanisms have been proposed to explain this phenomenon and its relationship to metabolic imbalances (Cani et al., 2019). These include endocrine regulation of food intake (Goswami et al., 2018; Lin et al., 2012), additional energy liberated by the microbiota from dietary fibers (Turnbaugh et al., 2006), alterations in bile acid profiles (Sayin et al., 2013; Yao et al., 2018), inflammatory responses induced by some members of the microbiota (Caesar et al., 2015) and induction of thermogenesis in adipose tissue (Krisko et al., 2020; Li et al., 2019, 2021). However, given the complexity of a complete microbiota and its interactions with the host, validating any of these theories and identifying causal relationships remains a major experimental challenge (Harley and Karp, 2012; Walter et al., 2020).

Gnotobiotic mice, colonized with a simplified microbiota made up of defined species, have become a major tool to identify potential mechanisms of interaction between the microbiota and host (Koh and Bäckhed, 2020; Mallapaty, 2017; Steimle et al., 2021). Such approaches can generate a mechanistic understanding of how external factors (i.e. diet, infection) act on the different microbiota members individually and at a community level (Faith et al., 2014; Kovatcheva-Datchary et al., 2019). A widely used example, the OligoMM12, is a gnotobiotic consortium of 12 cultivable mouse-derived strains representing the major five bacterial phyla in the murine gut (Brugiroux et al., 2016). It is reproducible between facilities (Eberl et al., 2020) and extensive data now exists on the metabolism of individual species and their metabolic interactions with each other (Streidl et al., 2021; Weiss et al., 2021; Wotzka et al., 2019; Yilmaz et al., 2021). Understanding how, and to what extent, this gnotobiotic microbiota reconstitutes the metabolic phenotype of conventional mice is therefore of broad relevance for microbiota research.

Circadian variations in microbiota function adds an extra layer of complexity to metabolic interactions between the host and the microbiota. Circadian feeding is a major driver of microbiota composition (Thaiss et al., 2014; Zarrinpar et al., 2014). The luminal concentration of fermentation products such as short-chain fatty acids (SCFA) shows a dramatic circadian oscillation linked both to food intake and to intestinal motility (Tahara et al., 2018). Microbiota-derived molecules are known to influence host nutrient absorption (Wang et al., 2017) and host metabolic gene expression (Kuang et al., 2019; Thaiss et al., 2016). However, much of our current knowledge is derived from indirect calorimetry measurements made over a time period shorter than 24h (Bäckhed et al., 2004; Halatchev et al., 2019; Kübeck et al., 2016; Wostmann et al., 1968). Measurements of the same host-microbiota system, if taken at different timepoints in the circadian cycle of metabolism, could therefore be wrongly interpreted as qualitative shifts in microbiota function. Consequently, to understand the influence of the microbiota on host energy metabolism, it is key to quantify variation over the full circadian cycle.

A challenging aspect of addressing the influence of the OligoMM12 microbiota on host metabolism, is that long-term experiments require hygiene barrier conditions similar to those required to work with germ-free mice. In particular, standard metabolic cage systems do not permit maintenance of an axenic environment and moving mice between the open cages typically used in isolator systems where such animals are normally bred, to IVC-cage-like systems used for most metabolic cages, can be associated with stress and behavioral abnormalities (Rabasa and Dickson, 2016). We have therefore built an isolator-housed metabolic cage system. Based on the TSE PhenoMaster® system, we can monitor levels of O_2_, CO_2_ and hydrogen every 24min for up to 8 cages, across two separate isolators in parallel, while maintaining a strict hygienic barrier. This way, longitudinal monitoring of metabolism can be carried out over periods of several weeks in germ-free and gnotobiotic mice.

In this study, we applied this system to understand how well gnotobiotic microbiota replicate the influence of a complex microbiota on host metabolism. We compared the metabolic profile of germ-free (GF), gnotobiotic (OligoMM12) and conventionally raised mice (specific-opportunistic- pathogen free, SPF) fed *ad libitum* with standard chow, using isolator-based indirect calorimetry. Similar to what has been described before (Krisko et al., 2020; Kübeck et al., 2016), we found no significant differences in energy expenditure among GF and SPF mice. These results are in contrast to other work (Bäckhed et al., 2004; Levenson et al., 1969; Li et al., 2019; Wostmann et al., 1968), but the discrepancies can potentially be explained by the methods applied for normalizing energy-expenditure data. Germ-free and gnotobiotic mice exhibit extensive water retention in the cecal lumen which can contribute up to 10% of the total body weight. This water is metabolically inert but is included in the mass used for normalization in reports where a difference in energy expenditure is reported (Bäckhed et al., 2004; Levenson et al., 1969; Li et al., 2019; Wostmann et al., 1968). When accounting for cecal inert mass, no significant difference in energy expenditure can be found in either germ-free or the gnotobiotic OligoMM12 mouse line. By calculating consumed calories in food and waste calories in feces, we could replicate earlier findings that germ-free and gnotobiotic mice are less efficient at extracting calories from standard mouse chow. However, our calculations demonstrated that this is well-compensated by increased food intake such that all mice absorb a similar number of calories from food each day. Interestingly, despite indistinguishable energy expenditure, and indistinguishable energy absorption from food each day, OligoMM12 mice showed increased fat mass compared to both, germ-free and SPF mice. Consistent with alterations in energy storage patterns, their circadian patterns of respiratory exchange ratio (RER) and certain metabolites in liver and plasma, phenocopied SPF mice during the dark phase, but germ-free mice during the light phase. Our study indicates that a reductionist/synthetic microbiota can specifically recover microbiota function in the dark (active) phase, but not in the light (resting/fasting) phase of the circadian cycle. This represents a valuable tool for identifying critical microbiota species and functions needed to support healthy host metabolism throughout the day.

## RESULTS

To compare to published literature on germ-free and colonized mouse metabolism, we compared male, adult age-matched (12-14wks old) germ-free (GF), gnotobiotic (OligoMM12) and conventionally raised (SPF) mice, all bred and raised in flexible-film isolators and with a C57BL/6J genetic background. Indirect calorimetry measurements were carried out in flexible-film surgical isolators accommodating a TSE PhenoMaster® system (schematic view in Fig. 1A, picture Fig. 1B). Mice were adapted for between 24-36h to the single-housing condition inside isolator-based metabolic chambers before data collection. Variations on O_2_, CO_2_ and hydrogen, along with food and water consumption, were recorded every 24 min on each metabolic cage. We could confirm that germ-free mice maintain their germ-free status over at least 10 days of accommodation in these cages, via culture-dependent and culture-independent techniques (see Methods).

**Figure 1:**
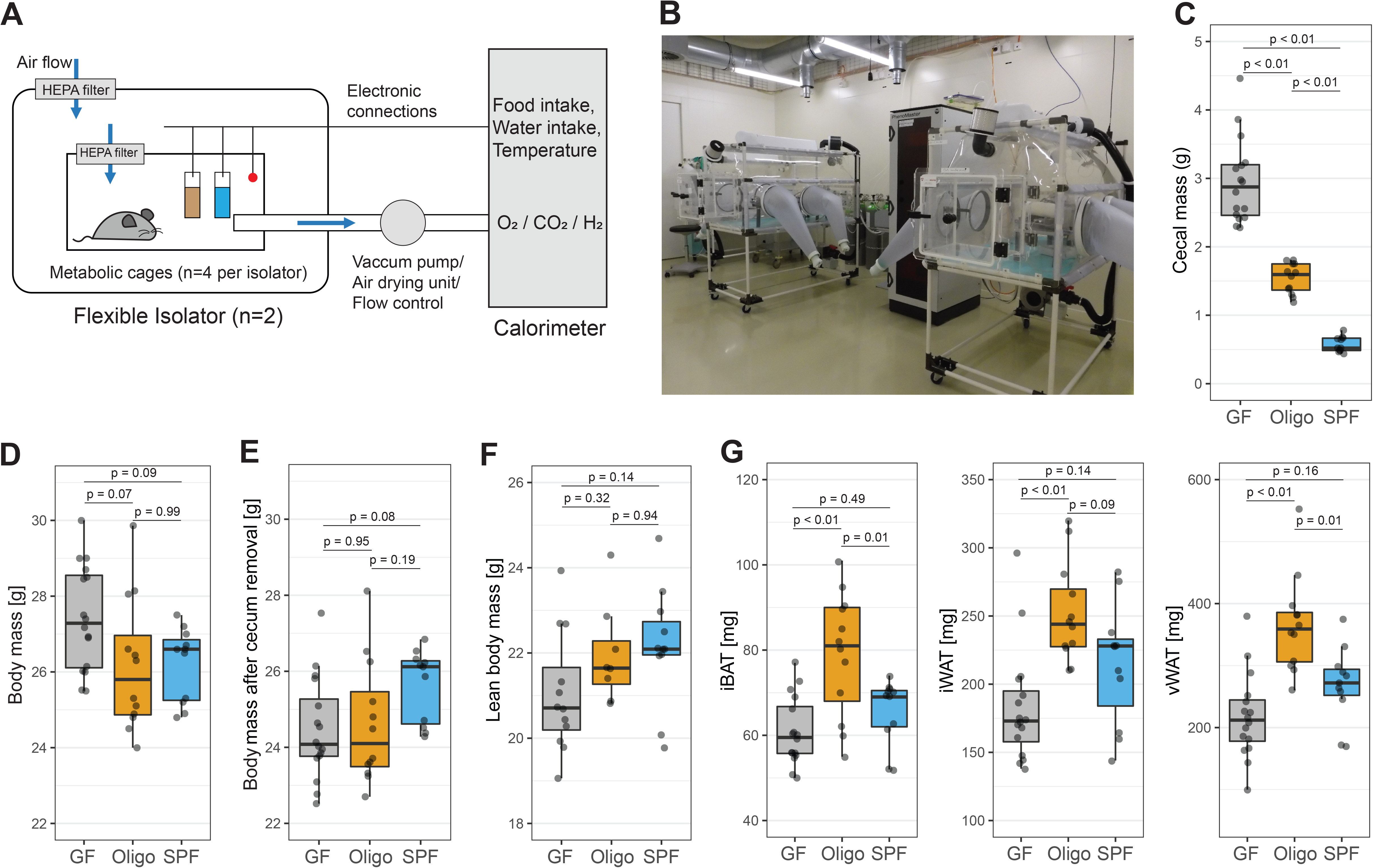
OligoMM12 mice have increase fat mass compared to GF mice and SPF C57B6/J mice. (A) Schematic representation of isolator-based indirect calorimetry system, with a TSE PhenoMaster® calorimeter connected to two flexible surgical isolators with four metabolic cages each. (B) Pictures of isolator-based indirect calorimetry system inside the facility. (C) Cecal mass (tissue including luminal content). (D) Total body mass at the end of the experiment and before cecum removal. (E) Total body mass after cecum removal. (F) Lean body mass acquired by EchoMRI before cecum removal (N of mice per group with EchoMRI and indirect calorimetry measurements: GF = 12, OligoMM12 = 8, SPF = 11). (G) Fat mass from interscapular brown adipose tissue (iBAT), inguinal white adipose tissue (iWAT) and visceral white adipose tissue (vWAT). Number of mice per group in all figures unless otherwise specified: GF = 16, OligoMM12 = 12, SPF = 11. p-values obtained by Tukey’s honest significance test.

### Body composition in GF, OligoMM12 and SPF mice

After 6-12 days of data recording mice were fasted for 4-5 hours and euthanized (approximately at ZT6 ± 1 hour), and body mass and body composition were measured. As cecal mass (cecal tissue plus its content) is affected by the colonization status (Wostmann et al., 1968), we first assessed the cecal mass in GF, OligoMM12 and SPF and its impact on body mass. We found that cecal mass was inversely correlated to the microbiota complexity, starting at approximately 0.5 g in SPF mice, increasing to around 1.5 g in OligoMM12 mice and reaching 3 g on average in GF mice (Fig.1C). Note that this represents around 10% of total body mass in GF mice (Suppl. Fig.1A), which translates into a trend to increased total body mass in GF mice (Fig, 1D). This trend was completely reverted after removal of the cecum from total mass (Fig. 1E).

Measurements of body composition in mice are often performed using EchoMRI, which yields data on lean, fat and water mass. As we observed that the cecum represented such a large and variable fraction of body mass, we compared EchoMRI read-outs of “lean” and “fat” body mass, before and after removal of the cecum (Suppl. Fig.1B-G). We found a strong correlation between the total lean mass measured by EchoMRI with and without the cecum (Suppl. Fig 1B) (Suppl. Fig.1C), i.e., cecum removal consistently reduced the lean mass readout by 5 to 10% (Suppl. Fig. 1D). Therefore, cecum removal has a consistent effect on lean mass across groups. For ease of comparison to published work, we decided to use lean mass obtained by EchoMRI *before dissection* for definitive energy expenditure calculations. We observed a trend in lean body mass with GF having a lower lean body mass than SPF mice, and OligoMM12 mice showing an intermediate phenotype (Fig. 1F). However, we were underpowered to detect a significant difference between groups, and we estimate that at least 22 mice per group would be needed to achieve statistical significance by one-way ANOVA if the current group differences are real – a number that was beyond the scope of the current study.

In contrast, EchoMRI fat mass measurements pre- and post-cecum dissection were poorly correlated in GF mice (Suppl. Fig. 1E) attributable to a variable scoring of cecal content as fat or water. In GF mice, cecum removal resulted in a decrease in EchoMRI fat mass readout of between 5 to 48% (Suppl. Fig. 1F) We also observed a shift towards *higher* fat mass readings in SPF mice after cecum removal (Suppl. Fig. 1F and Suppl. Fig. 1G); further highlighting the need for caution in interpreting EchoMRI readouts for fat mass in mice with major differences in intestinal colonization. Therefore, we proceeded to directly weigh the fat depots accessible to dissection (interscapular brown adipose tissue, iBAT; and, inguinal and visceral white adipose tissue, iWAT and vWAT). There was no significant difference between GF and SPF mice in size of the explored fat depots; however, OligoMM12 mice accumulated more fat in all explored depots than GF mice, including more iBAT and vWAT compared to SPF mice (Fig 1G).

### Energy metabolism and energy balance in GF, OligoMM12 and SPF mice

Body composition is determined by the quantity of calories absorbed from food, and whether these calories are directly expended or are stored. Energy expenditure was estimated using VO_2_ and VCO_2_ readouts (Meyer et al., 2015) and normalized as described before (Mina et al., 2018; Speakman, 2013; Tschop et al., 2011) using EchoMRI lean body mass and dissected fat mass.

In contrast to some previous reports (Bäckhed et al., 2004; Levenson, 1978; Li et al., 2019; Wostmann et al., 1968), but aligned with others (Krisko et al., 2020; Kübeck et al., 2016), we found no significant difference in daily energy expenditure (Fig. 2A and Fig. 2B) or VO_2_ (Suppl. Fig. 2A and Suppl. Fig. 2B) between GF, OligoMM12 and SPF mice after normalization using a regression model that included lean body mass and total dissected fat mass as predictive variables. This lack of difference was also observed when light and dark phases were analyzed separately (Fig. 2B and Suppl. Fig. 2B). “Classical” normalization procedures (dividing by mass) also showed no difference between groups when lean body mass, or “total body mass after cecum dissection” was used for normalization of energy expenditure (Fig. 2C and D) or VO_2_ (Suppl. Fig. 2C and D). Unsurprisingly, we did find a significant difference during the dark phase in energy expenditure (Fig. 2E) and VO_2_ (Suppl. Fig. 2E) between GF and SPF mice if “total body mass” (i.e., including the large inert cecum mass in germ-free and gnotobiotic mice) was used for normalization. Therefore, at least when comparing to the SPF microbiota used in this study, absence of a microbiota does not result in altered daily energy expenditure in metabolically active tissues.

**Figure 2:**
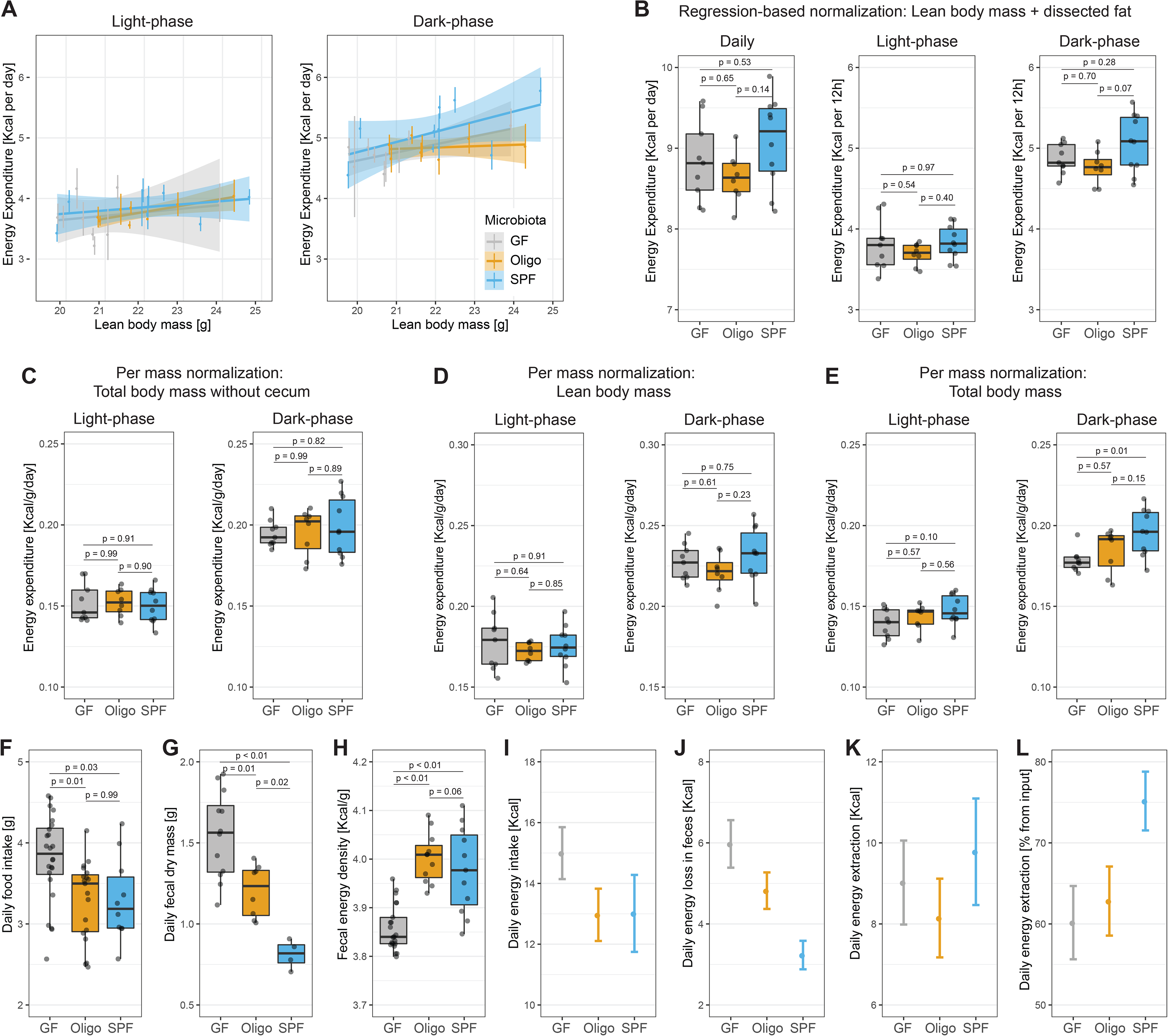
Energy metabolism in GF, OligoMM12 and SPF C57B6/J mice. (A) Linear regression of energy expenditure and lean body mass based on EchoMRI during light and dark phase. Each colored vertical line represents energy expenditure measurements during the experiment for one mouse. (B) Energy expenditure during 24h period, or during the 12h light or dark phase. Values represent area-under-curve normalized by regression-based analysis using lean body mass obtained by EchoMRI and dissected fat mass. (C, D, E) Energy expenditure values obtained by “classical” ratio-based normalization methods (dividing energy expenditure values per phase by mass). (C) Area-under-curve after normalization by total mass after cecal dissection. (D) Area-under-curve after normalization by lean body mass (EchoMRI). (E) Area-under-curve after normalization by total body mass before cecal dissection. (F) Average daily food intake per mouse. Mice represented in this figure include those that underwent long-term indirect calorimetry (Fig. 3) and mice that only contribute to daily fecal pellet quantification/bomb calorimetry. (N of mice per group: GF = 24, OligoMM12 = 19, SPF = 10) (G) Dry fecal output per mouse collected during a 24h period. (N of mice per group: GF = 12, OligoMM12 = 8, SPF = 4) (H) Energy content of dry fecal output by bomb calorimetry. (N of mice per group: GF = 21, OligoMM12 = 11, SPF = 11). (I, J, K, L) Estimation energy metabolism parameters. Number represented estimate mean value ± 1.96*combined standard uncertainty from measurements used for calculations. (I) Estimated daily energy input (food intake* 3.94 Kcal/g). (J) Estimated daily energy excretion (daily fecal dry mass*fecal energy content). (K) Estimated daily energy extraction (daily energy input – daily energy excretion). (L) Estimated energy extraction from food as percentage of energy input ((daily energy input - daily energy excretion)/daily energy input*100). Note that calculations in L, N and M are per mouse and are not normalized to body mass. Number of mice per group in all figures unless otherwise specified: GF = 9, OligoMM12 = 8, SPF = 10. p-values obtained by Tukey’s honest significance test.

We next investigated increased calorie absorption from food by comparing the daily energy ingestion from food and calorie excretion in feces of GF, OligoMM12 and SPF mice. The difference between these two values estimates the absorbed calories. As reported previously (Wostmann et al., 1983), GF animals ingested on average between 10-20% more standard chow compared to OligoMM12 and SPF mice (Fig. 2F). Correspondingly, GF animals also excreted a much larger dry mass of feces, while OligoMM12 mice produced an intermediate fecal mass and SPF mice excreted the least (Fig. 2G).

Remarkably, energy density of dry feces was lower in GF mice (3.7 Kcal/g) compared to colonized mice (OligoMM12 and SPF, 4.0 Kcal/g); with the latter showing no difference among them (Fig. 2H). This gap between GF and mice with microbiota can likely be explained by the fact that although fecal bacteria improve energy release from food, a considerable fraction of that energy remains stored in the bacteria present in the feces. Assuming certain averaged parameters (dry mass of a bacterium = 2.26x10^-13^g / bacteria cell (Dennis and Bremer, 2008), density of bacteria in feces = 5x10^11^ bacteria cells / g of feces (Barlow et al., 2020), and energy stored in bacteria = 4.58 Kcal/g of dry bacteria mass (Popovic, 2019)); we estimated that the fecal microbiota of colonized mice can contribute approx. 0.52 Kcal/g of dry fecal mass – slightly more than the energy density difference between fecal energy density in colonized and germ-free mice.

We then used these values for food intake, fecal dry mass output, and fecal energy density to estimate energy absorbed from the feces. We found that the higher food consumption in GF mice (Fig. 2I) almost perfectly counterbalance their corresponding higher energy excretion in feces (Fig. 2J), such that all mice extract around 9 Kcal per day from their food (Fig. 2K). This is consistent with our measurements of daily energy expenditure by indirect calorimetry (Fig. 2B), although it fails to explain the observed adiposity in the OligoMM12 mice (Fig. 1G). Unexpectedly, the OligoMM12 efficiency of release of calories from chow remains similar between germ-free and OligoMM12 mice. Given that the gut content of both OligoMM12 and SPF mice is densely colonized, and the fecal energy density is similar; it should be noted that the lower amount of energy extracted by the OligoMM12 is not so much due to a poorer digestive capacity of the gnotobiotic gut microbes, but rather that compared to the SPF microbiota, the calories extracted by the OligoMM12 microbiota are either retained within the microbes or not converted into compounds that can be taken up by their host (Fig. 2L).

We therefore concluded that daily energy expenditure and daily energy absorption from food vary only within the range of experimental error intrinsic to indirect calorimetry experiments. At a fundamental level, food intake therefore seems to be well regulated by microbiota-released calories. Despite this, OligoMM12 mice have an elevated fat mass. It remains a distinct possibility that gain of fat mass depends on the cumulative effect of very small differences in energy intake and energy expenditure that are simply not resolvable in our system. An alternative explanation is that microbiota composition influences energy storage. In order to gain a deeper insight into underlying mechanisms we carried out a series of more detailed analyses of metabolism.

### Circadian changes in RER and microbiota-derived hydrogen and short-chain fatty acids (SCFA)

Respiratory exchange ratio (RER, the ratio of CO_2_ produced per O_2_ consumed) is widely used as an informative proxy for substrate utilization (i.e., glucose or fatty acids) for oxidation in tissues. We observed that GF mice have a lower RER compared to SPF mice in both light and dark phases, indicative of increased fat/decreased glucose metabolism in GF mice (Fig 3A). These changes in RER are not related to differences in feeding patterns as all mice have a similar food intake patterns during the periods in which their RERs differ the most (Fig. 3B).

**Figure 3:**
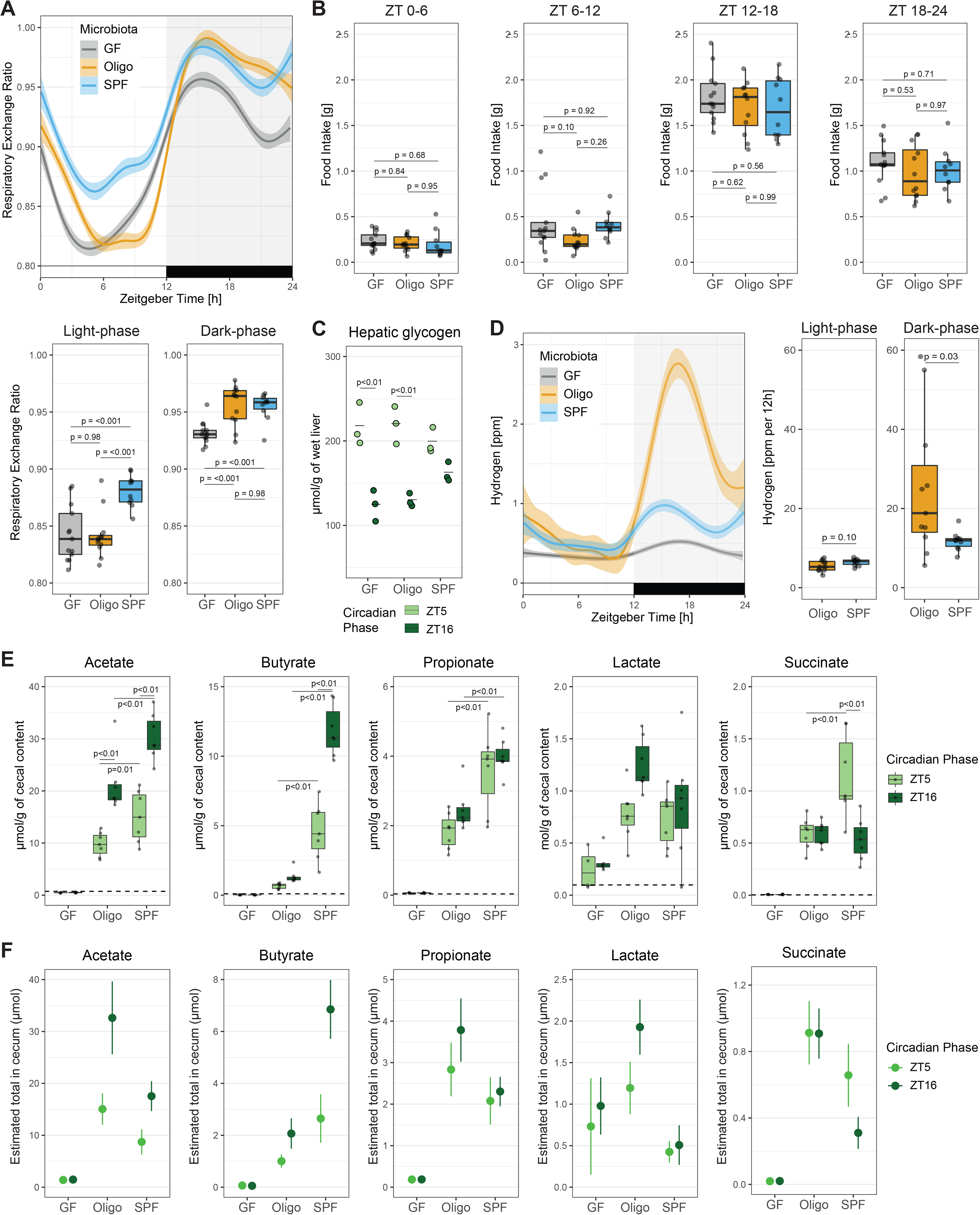
Circadian changes in Respiratory Exchange Ratio (RER), microbiota-derived hydrogen and short-chain fatty acids (SCFAs). (A) Comparison of circadian changes in RER among GF, OligoMM12 and SPF C57B6/J mice. RER curves obtained by smoothing function of data obtained every 24min per mouse over 10 days. Mean RER during the light phase (Zeitgeber 0-12) and dark phase (Zeitgeber 12-24). (B) Cumulative food intake during described ZT periods. Mice included in this analysis are those that underwent long-term indirect calorimetry, and they are a subset of the mice represented in Fig. 2F (C) Hepatic glycogen and triglyceride concentration in samples obtained at Zeitgeber 5 and 16 (N=3 per group). (D) Hydrogen production, curves obtained by smoothing function of data obtained every 24min per mouse. Area-under-curve after regression-based normalization by cecal mass during the light and dark phase (N of mice per group: OligoMM12 = 11, SPF = 10). (E) Concentration of short-chain fatty acids (acetate, butyrate, propionate) and intermediate metabolites (lactate, succinate) products in cecal content. Number of mice per group ZT5: GF = 4, OligoMM12 = 7, SPF = 7; ZT16: GF = 5, OligoMM12 = 7, SPF = 7. (F) Estimation total amount of short-chain fatty acids and intermediate metabolites by multiplying measured concentration values by the cecal mass of the group. Number represented estimate mean value ± combined standard uncertainty from measurements used for calculations. Number of mice per group in all figures unless otherwise specified: GF = 13, OligoMM12 = 12, SPF = 10. p-values obtained by Tukey’s honest significance test.

Differences in RER provided a clue that there could be differences in energy storage in mice with different microbiota status. Microbial fermentation products, including short-chain fatty acids (SCFA) and lactate, can be directly used as energy and carbon-sources by the murine host, and are generated by the microbiota via processes that liberate molecular hydrogen. We therefore quantified hepatic concentrations of glycogen, and cecal concentrations SCFA, at Zeitgeber 5 (ZT5, 5h into the light phase) and 16 (ZT16, 4h into the dark phase). Hydrogen was measured continuously during the circadian cycle.

Hepatic glycogen levels show a circadian rhythm, which usually peaks early during the transition between dark to light phase (ZT2-4), and drops to its minimum during the early hours of the dark phase (ZT14-16) in nocturnal rodents (Doi et al., 2010; Ishikawa and Shimazu, 1976). We found similar accumulation of hepatic glycogen in GF, OligoMM12 and SPF mice at ZT5; however, GF and OligoMM12 liver glycogen levels drop lower than SPF mice at ZT16 (Fig 3C). This differential pattern in GF/OligoMM12 compared to SPF mice may indicate that, although they can equally fill up hepatic glycogen storages at the end of the dark phase, GF and OligoMM12 deplete hepatic glycogen faster during the light phase.

Hydrogen, a byproduct of fiber fermentation by the microbiota, was also measured in the exhaust air of the metabolic cages. We found a clear circadian pattern in hydrogen production between OligoMM12 and SPF mice (Fig. 3D). Hydrogen levels in OligoMM12 and SPF mice decreased down to the limit of blank (GF level as reference) during the light phase, to later peak after food intake resumes during the dark phase. In addition, OligoMM12 mice showed a higher production of hydrogen than SPF mice during the dark phase even after regression-based normalization by cecal mass (Fig. 3D), i.e., the OligoMM12 microbiota produced hydrogen at a higher rate per cecal content mass than the SPF microbiota.

SCFA are the other major output of bacterial fermentation in the large intestine, as well as being key bioactive compounds produced by the large intestinal microbiota. SPF mice showed the highest cecal concentrations of acetate, butyrate, and propionate during both the light phase and dark phase, indicating efficient fermentation (Fig. 3E). Interestingly, OligoMM12 mice showed only 20-50% of the SCFA concentrations observed in SPF mice, but instead showed high production of lactate during the dark phase (Fig. 3E). In germ-free mice, all analyzed metabolites had levels below the limit of blank except for lactate, which could correspond to host-produced L-lactate (Zarrinpar et al., 2018) (our assay is not able to differentiate the enantiomers). As the total mass of cecum content is widely different among GF, OligoMM12 and SPF mice, we also estimated the total quantity of each compound in the cecal content by multiplying the concentration (Fig. 3E) by the cecal mass for each group (Fig. 1C) while propagating the uncertainty of each measurement.

This transformation has quite a major impact on how these data can be interpreted: when taking cecal mass into account, OligoMM12 mice have considerably higher levels of acetate during the light and dark phase and of propionate during the dark phase than SPF mice, while butyrate levels remain low. There is also an increased abundance of lactate and succinate in the OligoMM12 cecum content (Fig. 3F). Although we cannot directly link these microbial metabolites to the phenotype of the OligoMM12 mice, this underlines the major differences in metabolite profiles in the large intestine when comparing germ-free, gnotobiotic and SPF mice. High lactate production by the microbiome certainly warrants further study for potential metabolic effects on the host.

### Circadian changes in liver and plasma metabolites in GF, OligoMM12 and SPF mice

Finally, to increase our metabolic resolution, we applied UPLC-MS to perform untargeted metabolomics in the liver and plasma during the light (Zeitgeber 5) and dark phase (Zeitgeber 16) in GF, OligoMM12 and SPF mice. Correlating to what we observed in the RER during the light phase, GF and OligoMM12 cluster closely and are clearly separated from the SPF in the light phase of principal component analysis for both liver (Fig. 4A) and plasma samples (Fig. 4B). However, only minor shifts towards the SPF liver metabolome are seen during the dark-phase for OligoMM12 liver. This increased separation of the liver metabolome between germ-free and OligoMM12 mice during the dark-phase, is more apparent when SPF mice are excluded from the analysis (Suppl. Fig. 3A). Therefore, although RER and glycogen levels clearly show germ-free like patterns during the light-cycle and SPF-like patterns during the dark-phase, the underlying metabolome shifts attributable to the microbiome in OilgoMM12 mice are subtle, and generally closer to germ-free signatures than to SPF signatures in both liver (Fig. 4A) or plasma samples (Fig. 4B).

**Figure 4.**
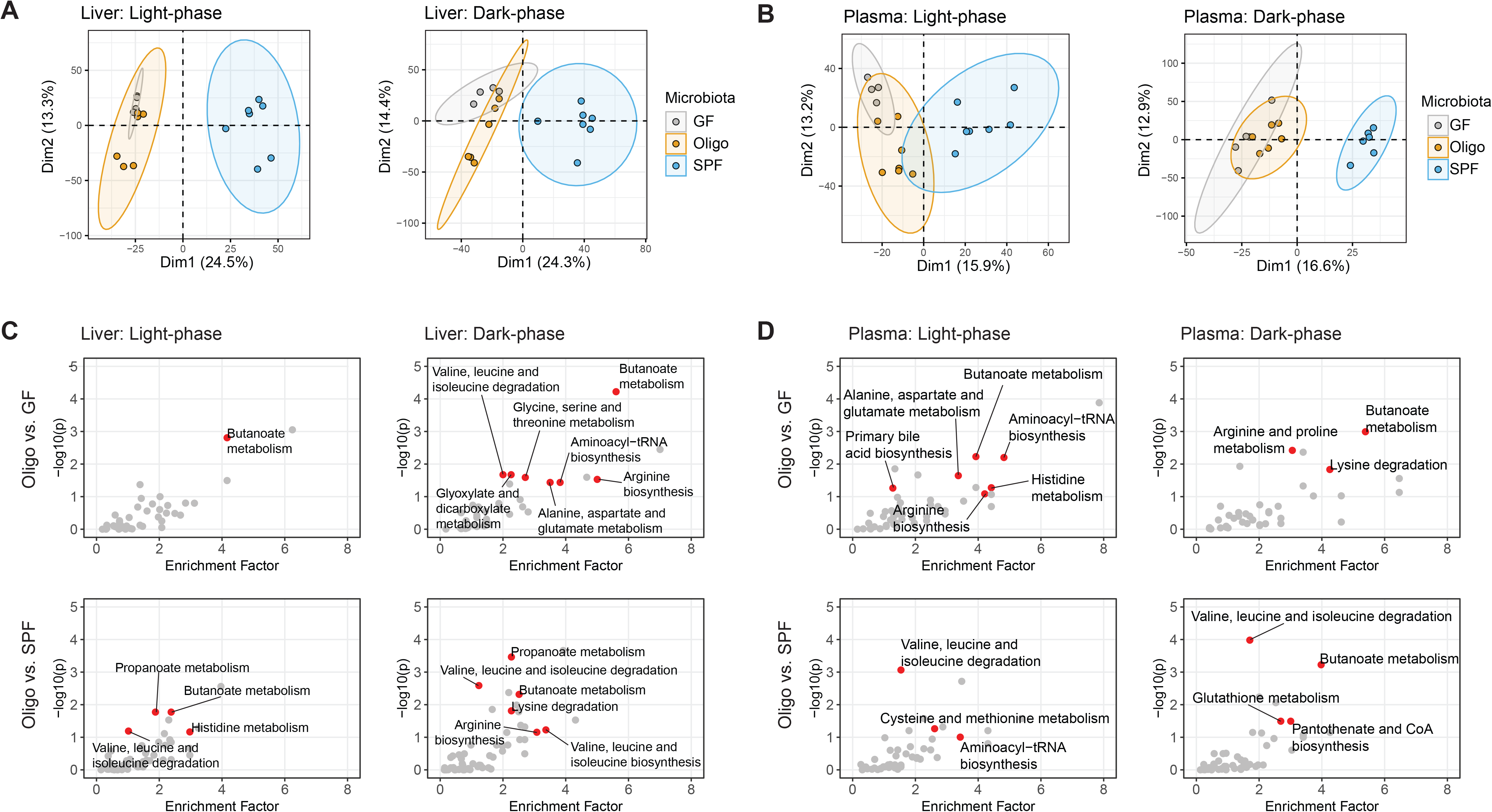
Metabolic profile comparison of GF, OligoMM12 and SPF C57B6/J mice by UPLC/MS. (A and B) Principal component analysis of metabolites identified by untargeted UPLC/MS during the light phase (Zeitgeber 5) and dark phase (Zeitgeber 16) in (A) liver and (B) plasma. (C and D) Metabolic pathways identified in the KEGG PATHWAY database; red dots represent pathways containing compounds differentially enriched in (*top*) OligoMM12 vs. GF and (*bottom*) OligoMM12 vs. SPF comparisons. Samples obtained during the light phase (Zeitgeber 5) and dark phase (Zeitgeber 16) in (C) liver and (D) plasma. Number of mice per group: Liver ZT5: GF = 4, OligoMM12 = 6, SPF = 7; ZT16: GF = 4, OligoMM12 = 6, SPF = 7 / Plasma: ZT5: GF = 4, OligoMM12 = 7, SPF = 7; ZT16: GF = 5, OligoMM12 = 6, SPF = 6.

We used the package MetaboAnalystR (Chong and Xia, 2018) to identify putative compounds that are significantly different in pair comparisons between OligoMM12 mice and their GF and SPF counterparts by untargeted peak extraction. These were then mapped onto metabolic pathways using the KEGG database. We found several pathways differentially enriched when OligoMM12 mice were compared to GF or SPF counterparts during the light and dark phase in liver (Fig. 4C) and plasma (Fig. 4D), including amino acid, bile acids, and fatty acid metabolism. Additionally, we selected compounds that belong to these differentially enriched pathways or have been previously identified to have circadian changes in obese patients (Nowak, 2021), confirmed their structure using chemical standards, and performed a targeted peak extraction for a more precise comparison among groups (Suppl. Table 1). We observed that for many of these metabolites the OligoMM12 microbiota produce an intermediate phenotype between GF and SPF mice, e.g., a subset of bile acids and amino acids, in liver (Suppl. Fig 4A) and plasma (Suppl. Fig 4B).

## DISCUSSION

Since the early days of nutritional studies, there has been a clear interest to understand the role of microbiota in host morphology, physiology and nutrition (Gordon and Pesti, 1971; Levenson, 1978). Pioneering work comparing germ-free rats with conventionally raised counterparts already described differences in food intake, energy extraction from diet and energy expenditure by indirect calorimetry (Levenson et al., 1969; Wostmann et al., 1983). More recently, researchers have explored the effect of specific complex microbiota communities and how they influence energy metabolism and body composition in the host (Ridaura et al., 2013; Suárez-Zamorano et al., 2015; Turnbaugh et al., 2006). Here we extend and clarify some of these observations via use of a well-established gnotobiotic mouse model consisting of 12 cultivable microbiota strains.

By carefully checking the validity of different measurement types, we found no significant difference in lean body mass among germ-free (GF), gnotobiotic (OligoMM12) and conventionally raised (SPF) mice. Interestingly, there was a significant increase in fat depots in OligoMM12 mice compared to GF and SPF animals. Previous studies diverged on the effect of microbiota on fat mass accumulation during conventional/low-fat diet feeding; reporting either increased fat mass in SPF mice (Bäckhed et al., 2004) or no difference compared to GF mice (Kübeck et al., 2016). However, it should be noted that there can be huge differences between SPF microbiota within and between animal facilities. GF mice transplanted with microbiota derived from obese donors accumulated more fat mass compared to those transplanted with microbiota derived from lean donors (Halatchev et al., 2019; Ridaura et al., 2013; Turnbaugh et al., 2006), with correlates identified to individual species/strain abundance (Woting et al., 2014, 2015). SPF microbiota matching more closely to those from obese donors could therefore be expected to give differing results to ours. In contrast, minimal microbiota communities such as the OligoMM12 can be perfectly replicated across sites (Eberl et al., 2020), and can help to clarify the complex processes linking microbiota and host metabolism (Becker et al., 2011). Further exploration of the metabolic effects of the OligoMM12 microbiota community, and extended versions thereof, has potential to clarify if specific strains, species or functional classes (Schmidt et al., 2018) are sufficient and necessary to drive the development of increased fat depots in these mice.

We further observed no significant difference in energy expenditure in GF, OligoMM12 and SPF. This was critically dependent on the mass normalization procedure applied. Normalization of mass-dependent variables by a per-mass (or allometric transformation) ratio has been recognized as a common source of controversy (Packard and Boardman, 1999; Tanner, 1949; White and Seymour, 2005), especially with large changes in body mass composition (Butler and Kozak, 2010; Kaiyala and Schwartz, 2011), and there have been several publications calling for the use of better statistical methods (Arch et al., 2006; Fernández-Verdejo et al., 2019; Tschop et al., 2011). Water- and indigestible solute retention in the cecum lumen of germ-free and gnotobiotic mice can contribute up to 10% of the total body mass and should be considered metabolically inert. It is therefore unsurprising that when the cecal content mass is very different among groups, using total body mass for normalization introduces a considerable bias in normalized energy expenditure estimation. Interestingly, it was long-ago observed that surgical removal of the cecum equalized the oxygen consumption between germ-free and conventional rats; as well as other measurements normalized by total body mass (Wostmann et al., 1968). With normalization using linear regression models based on lean-mass and fat-mass (Mina et al., 2018), we and others found no significant differences in energy expenditure by indirect calorimetry between GF and SPF mice under standard chow diet conditions (Krisko et al., 2020; Kübeck et al., 2016; Li et al., 2021).

An additional important confounder that we encountered was high variability of fat mass readouts obtained by EchoMRI when comparing mice with major differences in intestinal colonization levels. This could be attributed to variable calling of the fluid-filled ceca of gnotobiotic animals as either fat or water, compared with more accurate calling in conventional mice, revealing an important limitation of these systems. Consequently, physically dissected fat mass provided a more accurate read-out in all of these studies.

We are also keen to point out the more general limitations of our observations: only one gnotobiotic microbiota and one SPF microbiota were analyzed, and our conclusions pertain exclusively to these. We in no way exclude the possibility that some microbiota constituents or conformations *can* influence host energy expenditure (Halatchev et al., 2019) and/or body composition (Ridaura et al., 2013; von Schwartzenberg et al., 2021; Turnbaugh et al., 2006). In addition, it should be noted that indirect calorimetry is an inherently noisy data type, and small differences in daily energy expenditure are impossible to resolve via this technique (Corrigan et al., 2020; Fernández-Verdejo et al., 2019).

Nevertheless, the lack of measurable difference in energy expenditure between GF, OligoMM12 and SPF mice is aligned with our finding that the amount of energy obtained by *ad libitum* food intake was also remarkably similar among the groups. GF mice seem to accurately compensate the lower capacity of energy extraction from diet by increasing food intake. While this seems generally to be in agreement with models that described the regulation of appetite (and therefore energy intake) by the basal energy requirement of the individual (MacLean et al., 2017; Stubbs et al., 2018), it remains surprising given the discrepancy in the types of substrates available for oxidative metabolism in colonized and germ-free mice, revealed by RER differences. Although germ-free mice have a longer total gastrointestinal transit time than SPF mice (Touw et al., 2017), very little calorie absorption from food can occur after ingested food reaches the cecum of a germ-free mouse, whilst an SPF mouse will release usable energy from their food via microbial fermentation for several more hours in the cecum and colon, generating a major time-difference in the absorption of calories after eating in germ-free and SPF animals. This compensation seems also to function in mice colonized with the OligoMM12 microbiota, where despite robust microbial fermentation (read out as hydrogen and fermentation product production) and identical fecal energy density to SPF mice, energy recovery from ingested food is poor due to the volume of feces shed. A clear conclusion from these observations is that microbiota-dependent changes in metabolic substrates, and timing of calorie absorption, are well integrated in the murine central regulation of appetite over the course of a day (Fetissov, 2017).

Despite this broadly successful regulation of food intake and energy expenditure, at the molecular level, major differences were observed between the mice with different microbiota. First, OligoMM12 mice displayed an RER at the GF level during the light phase (when mice typically sleep and fast) but raised up to SPF levels during the dark phase (i.e., when mice are active and eating). It therefore appears that the OligoMM12 microbiota better recapitulates the microbiome effects on the host energy substrate use during the dark (active) phase when food-derived carbohydrates are abundant in the large intestine, but not in the light (sleeping) phase when mainly host-derived carbon sources are available in the large intestine. This potentially correlates with the SCFA concentrations observed in the cecum content of the OligoMM12, which was associated with a predominance of succinate and lactate, at the expense of propionate and butyrate. In complex microbiotas, lactate is typically further metabolized to butyrate by specific firmicutes (Belenguer et al., 2011; Duncan et al., 2004; Flint et al., 2015), which may be lacking or insufficiently abundant in the OligoMM12 mice. As lactate can inhibit lipolysis in adipocytes (Cai et al., 2008; Liu et al., 2009), this raises an interesting theme for follow-up studies to define the role of microbiota-derived lactate in host metabolism. In line with the RER data, we also observed that the liver and plasma metabolite profiles of OligoMM12 mice clustered closer to GF mice than to SPF mice. Although a small shift in the liver metabolome could be observed in the OligoMM liver during the dark cycle, this clearly demonstrates major metabolic effects of a complete microbiota that are not reconstituted by the OligoMM12 strains. In addition, certain amino acids were differentially represented between OligoMM12 and GF or SPF mice, as it has been described previously (Claus et al., 2008; Mardinoglu et al., 2015). Interestingly, OligoMM12 had a bile acid profile closer to GF than SPF mice, for example showing GF-levels of hepatic β-murocholic acid and taurine- β-murocholic acid, the predominant bile acid in the liver of GF mice (Sayin et al., 2013). Follow up studies with manipulation of the OligoMM12 microbiota or metabolic interventions are a promising tool to pull apart the circadian effects on RER, the influence of an unusual fermentation product profile, and other more subtle metabolic changes on overall metabolic health of the murine host.

In conclusion, our study showed that isolator-based indirect calorimetry is possible and allows detailed analysis of the metabolism of germ-free and gnotobiotic mice in real-time. Data generated with this system demonstrated that microbiota-released calories are well integrated in host energy balance, and that daily energy expenditure was not significantly influenced by microbiota composition in our mice. Nevertheless, mice colonized with the OligoMM12 gnotobiotic microbiota accumulated more fat mass and display a GF-like RER during the light phase but an SPF-like RER during the dark phase, indicative of altered metabolic substrate usage and energy storage. Correspondingly, the liver metabolome of mice colonized with the OligoMM12 showed alterations in bile acid, fatty acid and amino acid metabolism, despite overall clustering with the GF liver metabolome. This reveals the potential for gnotobiotic microbiota communities to investigate the mechanisms underlying the influence of microbiota on host metabolic health. As microbial dysbiosis is associated with a range of human diseases, circadian analysis of energy balance represents a crucial tool in the mining of microbiome data for therapeutic and diagnostic purposes.

## METHODS

### Animals

We used C57B6/J male mice aged between 12-14 weeks. We compare germ-free mice (GF), with a 12-strain gnotobiotic microbiota (Brugiroux et al., 2016) (OligoMM12) and specific-pathogen free mice (SPF). GF and OligoMM12 mouse lines are bred and maintained in open-top cages within flexible-film isolators, supplied with HEPA-filtered air, and autoclaved food and water ad libitum. As we are aware that housing conditions may influence behavior and potentially metabolism, we also bred and maintained a SPF colony under identical conditions inside a flexible-film isolator specifically for this study, such that all mice experienced identical living conditions, food, and water. Mice were adapted for between 24-36h after transfer from the breeding isolators to the isolator- based metabolic chambers. For long term experiments, mice were periodically re-housed in couples for short periods of times to avoid stress of extended single-housing conditions. In all cases, animals were maintained with standard chow (diet 3807, Kliba-Nafag, Switzerland) and autoclaved water. Germ-free status was confirmed at the end of the long-term experiments by culturing cecal content in sterile BHIS media in aerobic and anaerobic conditions for a week. In addition, cecal content was frozen at -20°C for a week, then stained with SYBR Gold and assessed by bacterial flow cytometry (Moor et al., 2016) using similarly processed SPF mice cecal content as positive control for the presence of bacteria. All experiments were conducted in accordance with the ethical permission of the Zürich Cantonal Authority.

### Indirect calorimetry

The isolator-housed TSE PhenoMaster® system allows instantaneous measurements of oxygen, carbon dioxide and hydrogen levels as well as total feed and water consumption while keeping a strict hygiene level of control. The metabolic isolator system consists of an adapted set of two flexible-film surgical isolators, each of them housing four metabolic cages from the TSE PhenoMaster® system (TSE Systems, Germany). Room air is pulled into the isolator by a vacuum pump passing through a double set of HEPA filters. Then, each cage is connected via a second HEPA filter through the back wall of the isolator to the CaloSys setup, which pulls sterile air from the isolator into the cages using negative pressure. Air coming from the cages is dehumidified at 4°C and sequentially passed by a Sensepoint XCD Hydrogen gas analyzer (Honeywell Analytics, Hegnau, Switzerland) and standard oxygen and carbon dioxide censors provided in the TSE PhenoMaster® system. A two-point calibration of all analyzers using reference gases was performed within 24 h before each animal experiment. Data was recorded using a customized version of the TSE PhenoMaster® software modified to integrate hydrogen measurements.

For indirect calorimetry measurements, the animals were transported in pre-autoclaved, sealed transport cages from the breeding isolators into the metabolic isolator system. Mice were single housed and adapted for between 24-36h before starting recording measurements to ensure proper access to food and water as well as account for initial exploratory behavior. Mice were kept up to 10 days at a stable temperature (21-22°C) with *ad libitum* availability of standard chow and water. The days were divided into a dark and light period of 12 hours each. In this study, we kept the air flow of 0.4 L/min and recorded individual cage data (gases production and food/water consumption) every 24min (time set per cage for measurement stabilization 2.5min). In long experiments, mice were periodically pair-housed for 24h to prevent stress due to prolonged single housing.

### Body composition measurements

At the end of the experiment, mice were fasted for 4 hours (Zeitgeber 1 till 5) before for body composition measurements. We used magnetic-resonance whole-body composition analyzer (EchoMRI, Houston, USA) to analyze mice body composition (lean and fat mass). Then, mice were euthanized using CO_2_ according to approved protocols. Total body mass was obtained by weighing the full carcass and cecum was dissected and weighed by one investigator (DH). For a set of mice, we remeasured body composition by EchoMRI after cecum removal. Finally, fat depots were dissected from all mice by one investigator (W.S.) that was blinded to the hygiene status of the mice. Interscapular brown adipose tissue (iBAT), inguinal white adipose tissue (iWAT) and visceral white adipose tissue (vWAT) were sampled and weighted.

### Food intake, fecal samples and bomb calorimetry

Daily food intake was obtained as the mean value of food intake recorded by the TSE PhenoMaster® system during the course of the experiment. In addition to the mice reported in the indirect calorimetry experiments, we also collected food intake data from a set of selected experiments in which we collected fecal pellets produced during 24h. For daily fecal excretion measurements, we cleaned up the bedding in the cage and replaced it for a clean and reduced amount of bedding. After 24h, we collected the mix of bedding and fecal pellets. Fecal pellets were manually collected from the bedding, transferred to 15ml tubes and stored at -20°C until bomb calorimetry. Before bomb calorimetry, fecal samples were freeze dried in a lyophilizer overnight (ALPHA 2-4 LDplus, Christ, Germany) and dry mass recorded. We used a C1 static jacket oxygen bomb calorimeter (IKA, Germany) to quantify the residual energy present in these dry fecal pellets, using approximately 0.2-0.5g of material. Energy content was normalized to grams of dry fecal pellets.

### Metabolomics by UPLC/MS

#### Sample obtention and preparation

Approximately at Zeitgeber 5 and 16, three mice of each group were euthanized, and liver and plasma samples collected. To minimize variations among mice, individual mice were euthanized with CO_2_ and sampled as fast as possible. Blood was obtained by cardiac puncture, collected in lithium heparin coated tubes, and kept on ice for further processing. Mice were perfused with PBS and liver samples were obtained by dissection of the lower right lobe, collected on an 2ml Eppendorf tube and flash frozen in liquid nitrogen. Finally, between 60-80 mg of cecal content was collected in a 2ml Eppendorf tube and flash frozen in liquid nitrogen. After samples all samples were obtained, blood samples were centrifuged 8000rcf for 5min, supernatant collected, and flash frozen in liquid nitrogen. Samples were kept at -80°C until preparation for UPLC/MS.

#### Short chain fatty acid quantification by UPLC/MS

Samples were first homogenized in 70%-isopropanol (1 mL per 10 mg sample), centrifuged. Supernatants were used for SCFA quantification using a protocol similar to previously described (Liebisch et al., 2019). Briefly, a 7-points calibration curve was prepared. Calibrators and samples were spiked with mixture of isotope-labeled internal standards, derivatized to 3- nitrophenylhydrazones, and the derivatization reaction was quenched by mixing with 0.1% formic acid. Four µL of the reaction mixture was then injected into a UPLC-MS system, [M-H]- peaks of the derivatized SCFAs were fragmented and characteristic MS2 peaks were used for quantification.

#### Untargeted UPLC/MS

Samples were thawed on ice. Serum samples were diluted with 90% methanol in water with a volumetric ratio of 1:7, incubated for 10 min on ice for allowing protein to precipitate. Liver samples were mixed with 75% methanol in water (2 mL/ 100 mg liver), homogenized using a TissueLyser (Qiagen, Germany) at 25 Hz for 5 min. The result mixtures were centrifuged at 15,800 g, 4 °C for 15 min. 100 µL of the supernatants were filtered with 0.2 µm reversed cellulose membrane filter and transferred to sample vials and used for UPLC/MS analysis with an ACQUITY UPLC BEH AMIDE column (1.7 µm, 2.1 × 150 mm, Waters). Another 400 µL of the supernatants were then lyophilized and resuspended in 80 µL 5% methanol in water, sonicated, filtered, and used for UPLC/MS analysis with an ACQUITY UPLC BEH C18 column (1.7 µm, 2.1 × 150 mm, Waters, RP column).

An ACQUITY UPLC system (I-Class, Waters, MA, USA) coupled with an Orbitrap Q-Exactive Plus mass spectrometer (Thermo Scientific, San Jose, CA) were used for UPLC/MS analysis. For the AMIDE column a flow rate of 400 µL/min was used with a binary mixture of solvent A (water with 0.1% formic acid) and solvent B (acetonitrile with 0.1% formic acid). The gradient starts from 1% of A, then gradually increases to 70% of A within 7 min. Then a 1% of A is kept for 3 min. The column was kept at 45 °C and the autosampler at 5° C.

For the RP column, the flow rate was set to 240 µL/min using a binary mixture of solvent A (water with 5 % methanol and 0.1 % formic acid) and solvent B (methanol with 0.1 % formic acid). The gradient starts from 95% of A, then gradually decreases to 5% of A within 10 min. A 100% solvent of B is kept for 2 min, then a 100% of A is kept for 2min to restore the gradient. The column was kept at 30 °C and the autosampler at 5 °C.

The MS was operated at a resolution of 140,000 at m/z = 200, with automatic gain control target of 2x10^5^ and maximum injection time was set to 100ms. The range of detection was set to m/z 50 to 750. Untargeted MS data was extracted from raw MS files by using XCMS (Smith et al., 2006) in R (v3.6.1), and then subject to pathway enrichment by using MetaboAnalystR (Chong and Xia, 2018).

#### Compound identification and targeted peak extraction

Chemical standards of selected compounds were diluted to 10 µg/mL and were analyzed using the UPLC/MS methods described before. Identification was done by comparing retention time and MS2 in liver/plasma samples with the chemical standards (Nowak, 2021). After confirming the chemical identities of the compounds, targeted peak extraction was done using Skyline (v21.1) (Adams et al., 2020).

### Data Analysis

#### Data quality control

To facilitate analysis across different experimental runs, all times were converted into Zeitgeber time (ZT; [h]), where 0-12 represents the light phase and 12-24 represents the dark phase. Any datapoint taken before the start of the first occurrence of ZT=0 was discarded. To account for faulty measurements caused by measurement imprecision, equipment malfunction or other disruptive events, datapoints were removed from the raw datasets according to criteria based on statistical and biological arguments. Food consumption values of 0.01 g during the 24min intervals were considered as measurement noise and discarded. Negative values for food and water consumption, as well as oxygen (dO_2_) and carbon dioxide (dCO_2_) differentials between the measurement chambers and the reference chamber were also considered as measurement noise and discarded. For the remaining subsets of measurements from the individual mice, we cleaned up outlier measurements in food and water intake by eliminating values greater than 75^th^ percentile + 1.5 times interquartile range. Potential sources for outlier measurements in food and water consumption observed included leaky water bottles and loss of food pellets during mice husbandry procedures. A similar approach was used to eliminate outliers from dO_2_ and dCO_2_ values below 25^th^ percentile - 1.5 times interquartile range. Potential sources for outlier measurements in gas differentials included inappropriate sealing of individual metabolic cages or clogging of pre-analyzer filters. Oxygen consumption (VO_2_) and CO_2_ production (VCO_2_) was calculated using dO_2_ and dCO_2_ and the Haldane transformation as described before (Arch et al., 2006). Energy expenditure was estimated from dO_2_ and dCO_2_ using Weir’s approximation (Weir, 1949). As one of the study objectives is to explore circadian patterns, if more than 20% of datapoints had to be removed from a particular day for a particular mouse, all other datapoints from that subset were discarded as well. After the cleanup process described above, the data from all different experiment runs were pooled together for further analysis. The above processes lead to a reduction in dataset size from 10472 to 9453 entries.

#### Statistical analysis

From the resulting dataset, energy expenditure over a certain period was calculated as the area under the curve (trapezoid interpolation) of instantaneous values obtained during the 24min measurements intervals. Food intake values calculated over a certain time are always cumulative. To compare different mice in the above variables, variations in body mass and composition between individuals need to be accounted for. As suggested in several publications (Fernández-Verdejo et al., 2019; Speakman, 2013; Tschop et al., 2011), this was done by regression-based analysis of covariance (ANCOVA). As such, a linear regression is performed on energy expenditure as a function of lean body mass and fat depots mass, with the microbiota group as a qualitative covariate. Then, each individual value is replaced by the sum of the corresponding residual and the energy expenditure predicted by the linear model using the average lean body and fat depot mass (calculated over all groups). Hydrogen production (difference in hydrogen concentration between the measurement chambers and the reference chamber) was adjusted in analogous fashion, using cecal mass (as a proxy for total gut microbiota mass) as a predictor.

For variables where the continuous evolution during the circadian cycle is of interest (RER, gross hydrogen production), values were averaged at each time point for each individual. A generalized additive model was used to fit a smooth line to these averages using a cubic penalized regression spline (using R function mgcv::gam with formula y ∼ s(x; bs = “cs”)).

For estimating derived variables (i.e., daily energy excretion) we used the R package “errors” (Ucar et al., 2019). This package links uncertainty metadata to quantity values (i.e., mean “daily fecal dry mass excretion”, mean “fecal energy content”) and this uncertainty is automatically propagated when calculating derived variables (i.e., “daily energy excretion” = “daily fecal dry mass excretion” x “fecal energy content”). Uncertainty is treated as coming from Gaussian and linear sources and propagates them using the first-order Taylor series method for propagation of uncertainty.

Hierarchical clustering and heatmap visualization were produced using the R package “pheatmap” using Pearson correlation as distance measure for clustering and Ward’s minimum variance method using an algorithm that includes Ward’s criterion (Murtagh and Legendre, 2014). For the Principal Component Analysis, we used the *prcomp* function which is present in built-in R stats and the R package “factoextra” for visualization.

All group comparisons were analyzed by ANOVA and Tukey’s honest significance test. For comparisons of metabolites identified by targeted peak extraction among groups, area values were log2 transformed before the statistical test.

## Resource availability

### Lead contact

Any further communication, including those related to resource sharing, may be directed to and fulfilled by the lead contact Emma Slack.

### Materials availability

This study did not generate new unique reagents.

### Data and code availability

Source data for Fig. 1, 2 and 3, and Suppl. Fig. 1 and 2 are available in the Supplementary Information. Source data for Fig. 4 and Suppl. Fig. 3 and the datasets and code used for all figures in this publication are made available in a curated data archive at ETH Zurich (https://www.research-collection.ethz.ch/handle/20.500.11850/501168) under the DOI https://doi.org/10.3929/ethz-b-000501168.

## ACKNOWLEDGEMENTS

We would like to thanks to Sven Nowok, Thomas Fehr, Andre Galhano and Susanne Freedrich for their support in the establishment of the gnotobiotic metabolic phenotype facility in the ETH Phenomic Center. Also, we thank Maria L. Balmer for her comments and suggestions for the manuscript. The gnotobiotic metabolic phenotype facility was initially funded by the FreeNovation grant (Novartis). E.S. and W-D.H. are supported by the NCCR Microbiomes, a research consortium financed by the SNF. E.S. acknowledges the support of the Swiss National Science Foundation (40B2-0_180953, 310030_185128), European Research Council Consolidator Grant, and Gebert Rüf Microbials (GR073_17) and the Botnar Research Centre for Child Health Multi- Invesitigator Project 2020. This project is part of “Zurich Exhalomics”, a flagship project of “Hochschulmedizin Zürich”.

## Author Contributions

Conceptualization, D.H., W-D.H., C.W. and E.S.; Methodology, D.H., J.L., W.S., R.Z. C.W. and E.S.; Formal Analysis, D.H., J.L, W.S., T.G. and M.A.; Investigation, D.H., J.L., W.S. and S.N.; Resources, B.S., R.Z., W-D.H., C.W. and E.S.; Writing – Original draft, D.H. and E.S.; Writing – Review and Editing D.H., J.L., W.S, T.G, M.A, S.N., A.J.M., B.S., R.Z., W-D.H., C.W and E.S.; Visualization, D.H., W-D.H., C.W. and E.S.; Supervision, W-D.H., C.W. and E.S.; Funding acquisition, W-D.H., C.W., A.J.M, and E.S.

## SUPPLEMENTAL INFORMATION TITLES AND LEGENDS

**Supplementary Figure 1:**
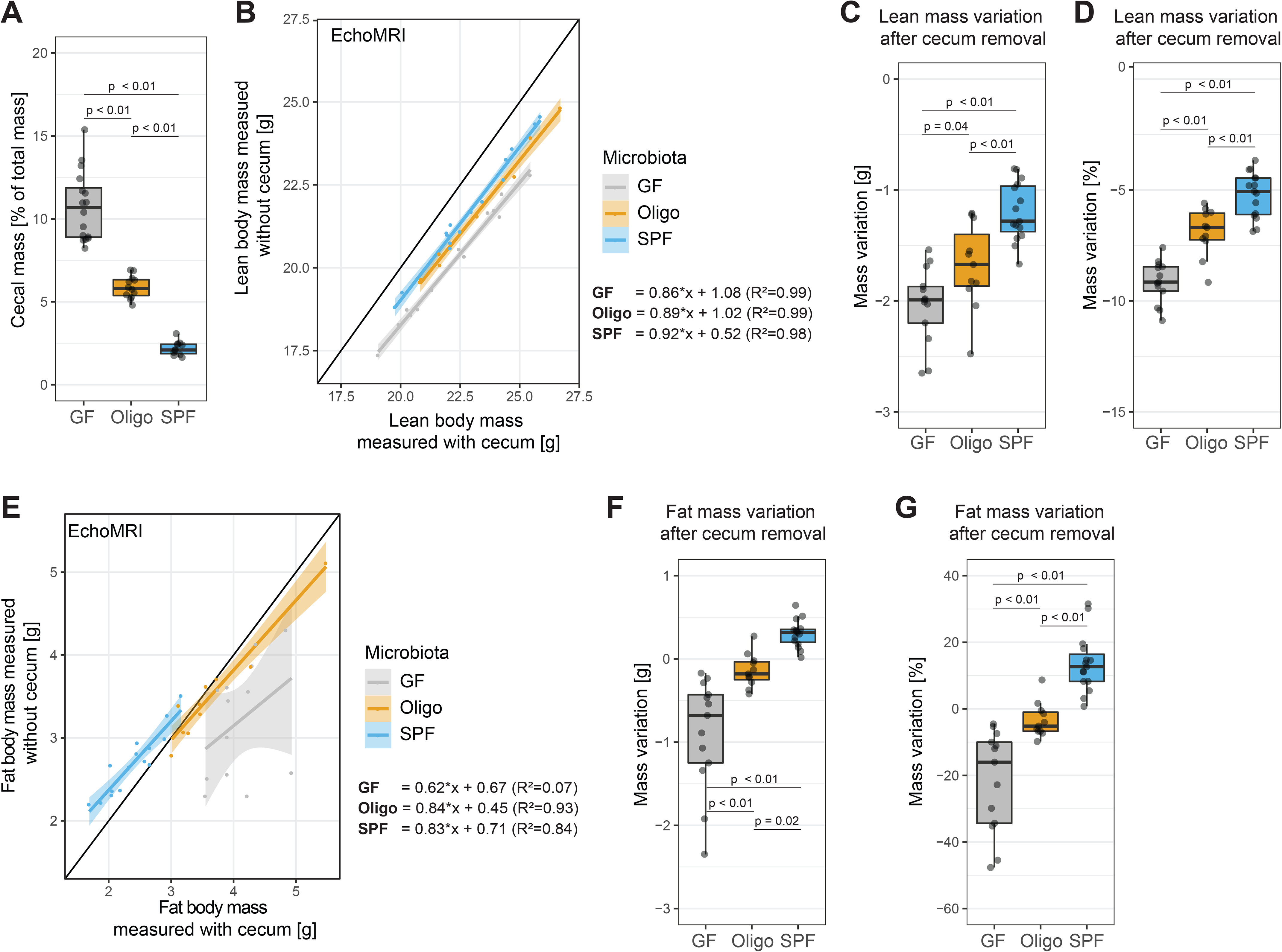
Cecal mass interferes with fat mass estimation by EchoMRI. (A) Cecal mass (tissue including luminal content) as percentage of total body mass (N of mice per group: GF = 16, OligoMM12 = 12, SPF = 11) (B) Lean body mass estimated by EchoMRI with and without cecum. Equations show simple linear regression for estimating lean mass without cecum based on lean mass with cecum; in brackets adjusted-R squared. (C) Lean mass variation after cecum removal. (D) Lean mass variation after cecum removal as percentage of lean mass before cecum removal. (E) Fat body mass estimated by EchoMRI with and without cecum. Equations show simple linear regression for estimating fat mass without cecum based on fat mass with cecum; in brackets adjusted-R squared. (F) Fat mass variation after cecum removal. (G) Fat mass variation after cecum removal as percentage of lean mass before cecum removal. Number of mice per group in all figures unless otherwise specified: GF = 13, OligoMM12 = 11, SPF = 15. p-values obtained by Tukey’s honest significance test.

**Supplementary Figure 2:**
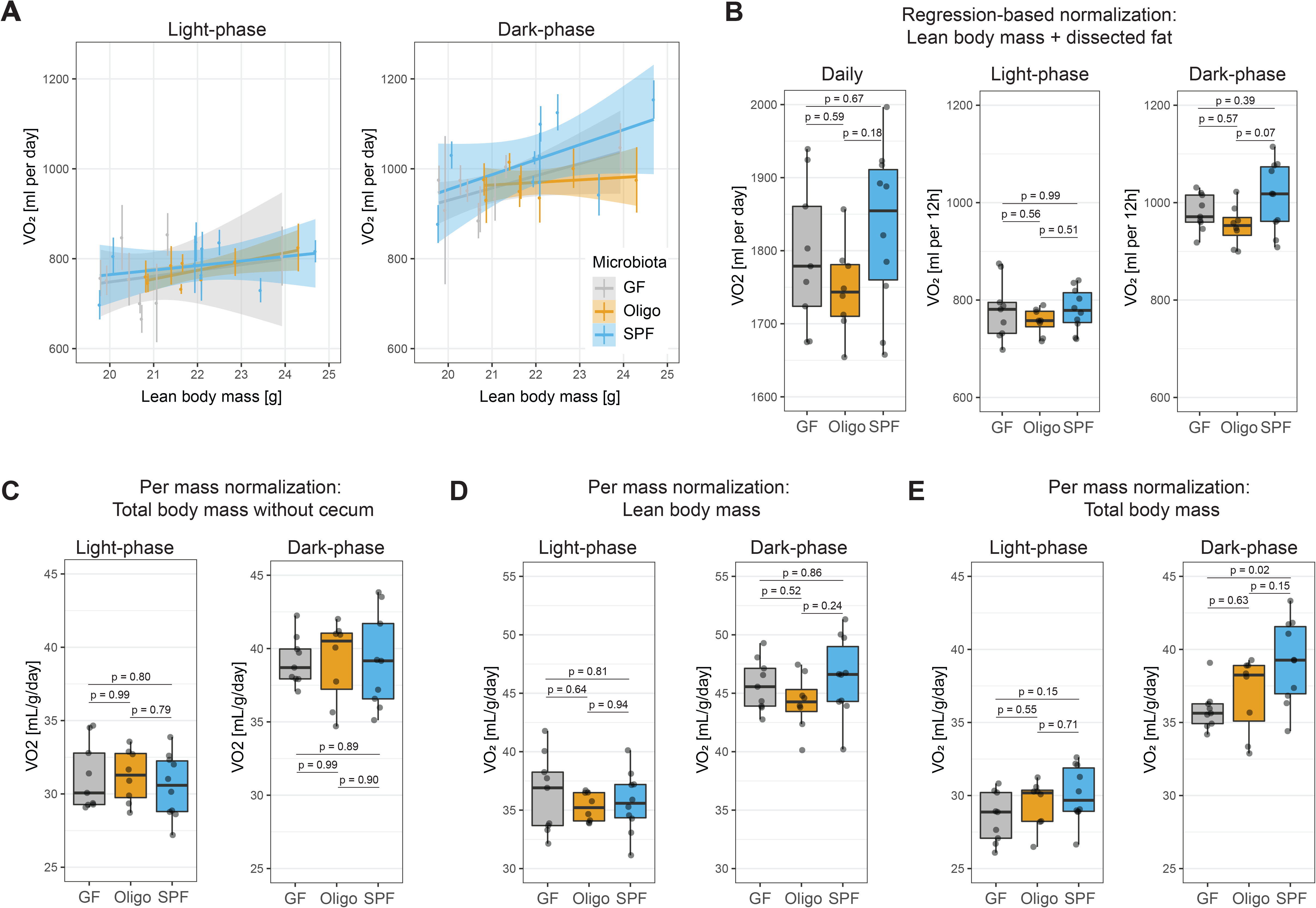
Cecal mass interferes with normalization of VO_2_. (A) Linear regression of VO_2_ and lean body mass (EchoMRI) during light and dark phase. Each colored vertical line represents energy expenditure measurements during the experiment per mouse. (B) VO_2_ during 24h period, or during the 12h light or dark phase. Values represent area-under-curve normalized by regression-based analysis using lean body mass obtained by EchoMRI and dissected fat mass (C, D, E) VO_2_ values obtained by “classical” ratio-based normalization methods (dividing energy expenditure values per phase by mass). (C) Area-under-curve after normalization by total mass after cecal dissection. (D) Area-under-curve after normalization by lean body mass. (E) Area-under-curve after normalization by total body mass before cecal dissection. Number of mice per group in all figures unless otherwise specified: GF = 9, OligoMM12 = 8, SPF = 10. p-values obtained by Tukey’s honest significance test.

**Supplementary Figure 3.**
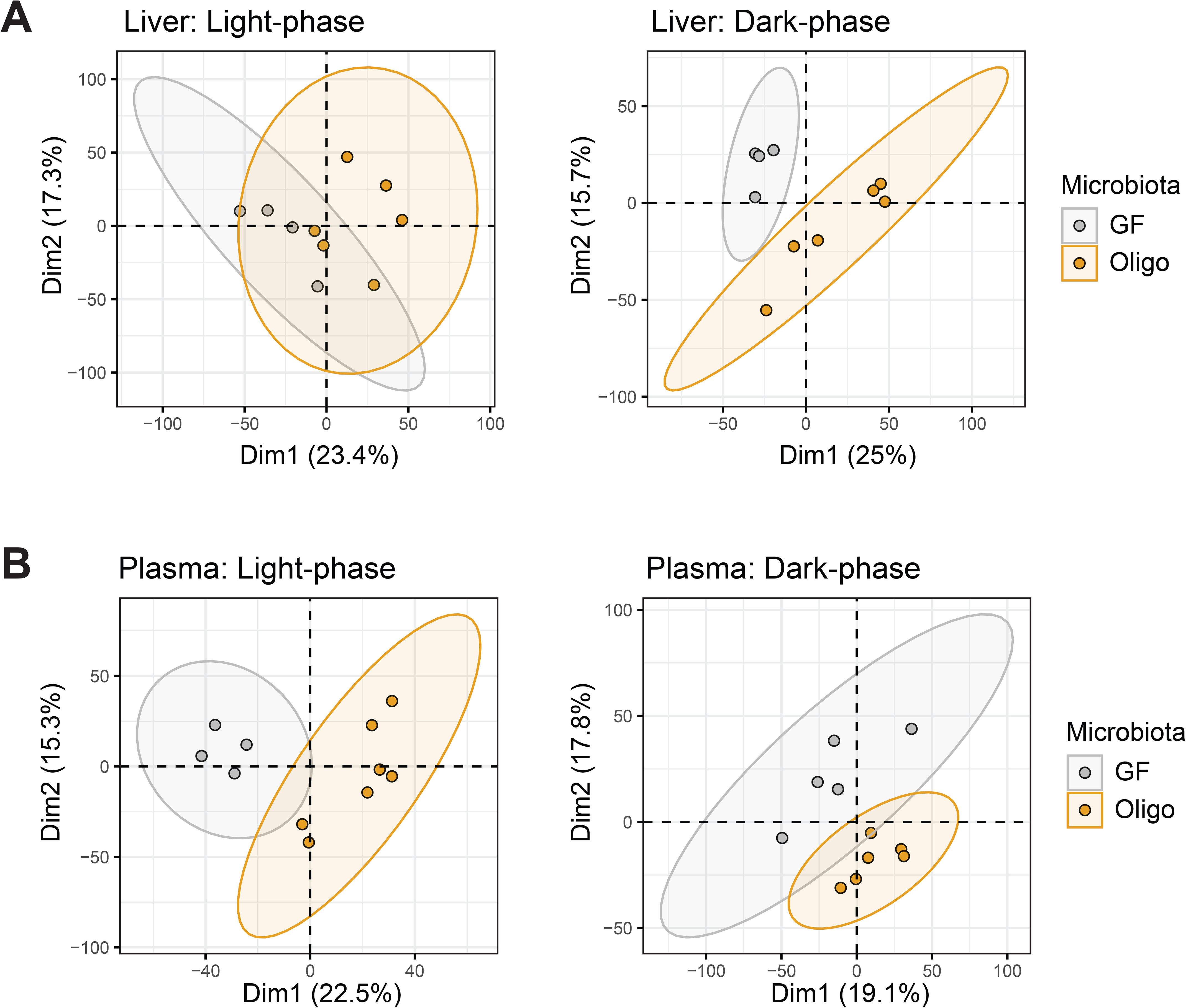
**Metabolic profile comparison of GF and OligoMM12 C57B6/J mice by UPLC/MS**. (A and B) Principal component analysis of metabolites identified by untargeted UPLC/MS during the light phase (Zeitgeber 5) and dark phase (Zeitgeber 16) in (A) liver and (B) plasma.

**Supplementary Figure 4.**
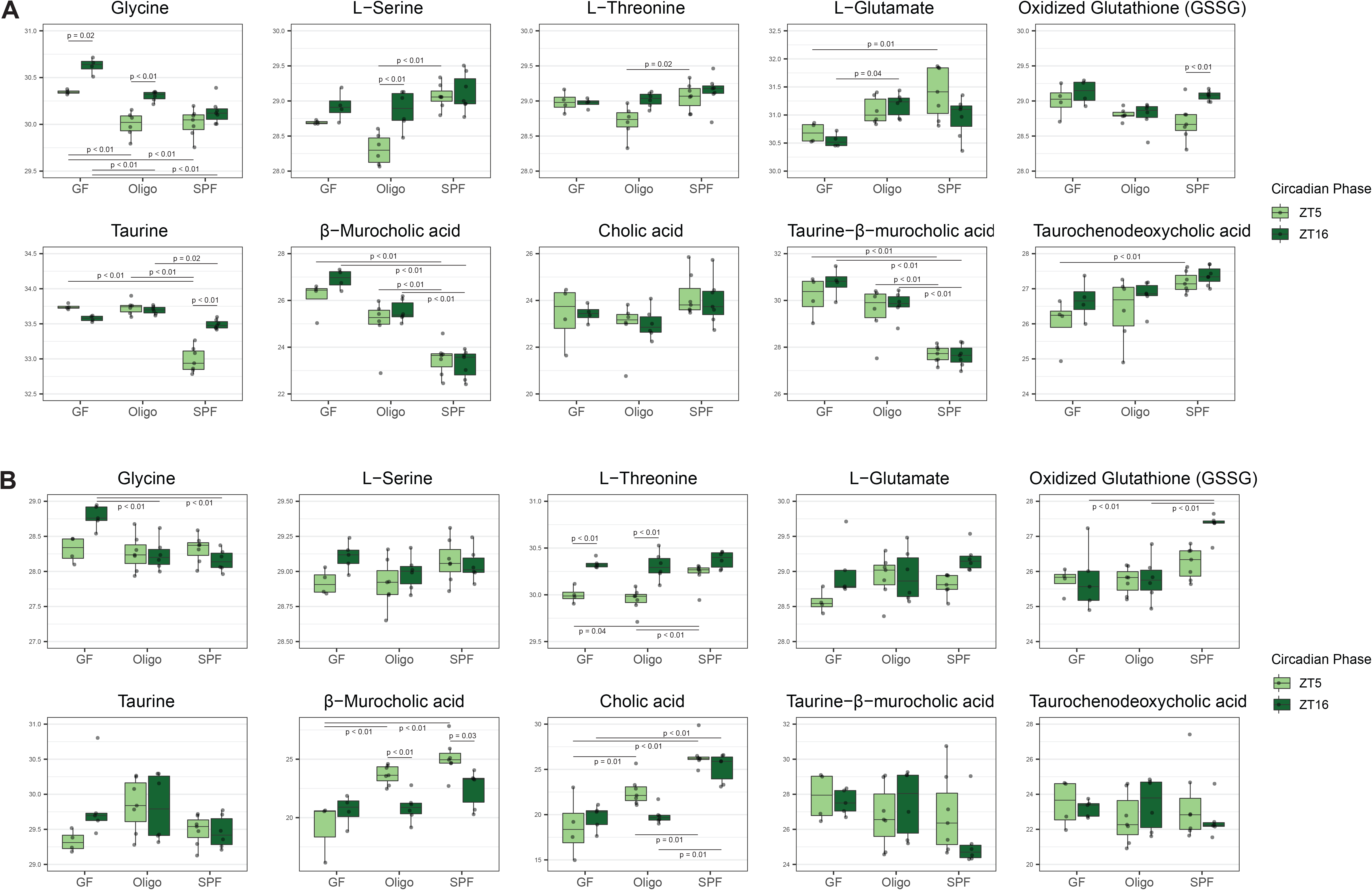
Metabolic profile comparison of GF, OligoMM12 and SPF C57B6/J mice by UPLC/MS. (A and B) Manually-curated list of compounds obtained by targeted peak extraction from differentially expressed pathways in (A) liver and (B) plasma samples. p-values obtained by Tukey’s honest significance test after log2 transformation of area value. Number of mice per group: Liver ZT5: GF = 4, OligoMM12 = 6, SPF = 7; ZT16: GF = 4, OligoMM12 = 6, SPF = 7 / Plasma: ZT5: GF = 4, OligoMM12 = 7, SPF = 7; ZT16: GF = 5, OligoMM12 = 6, SPF = 6.

## SUPPLEMENTARY TABLE

**Supplementary Table 1.**
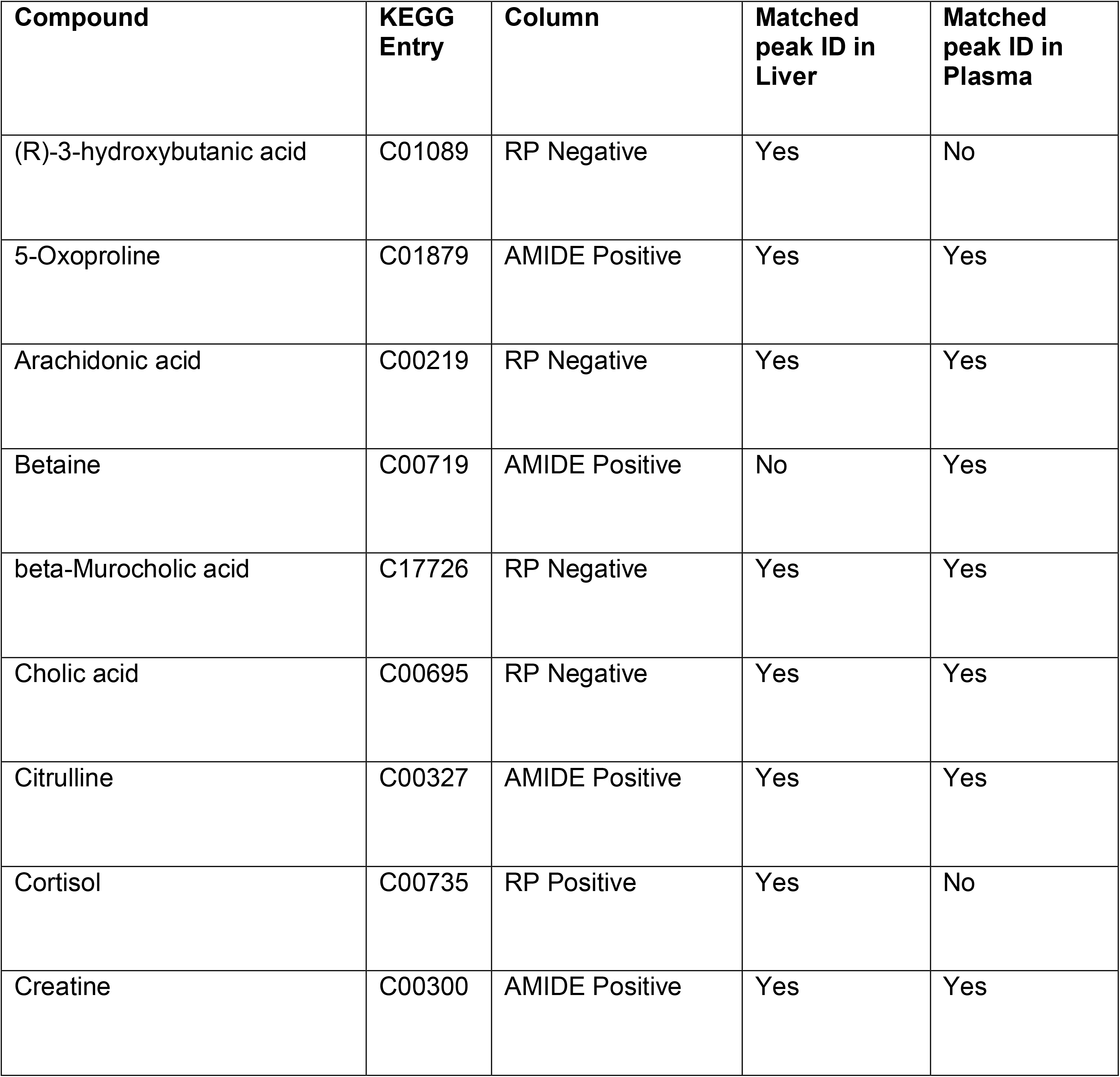
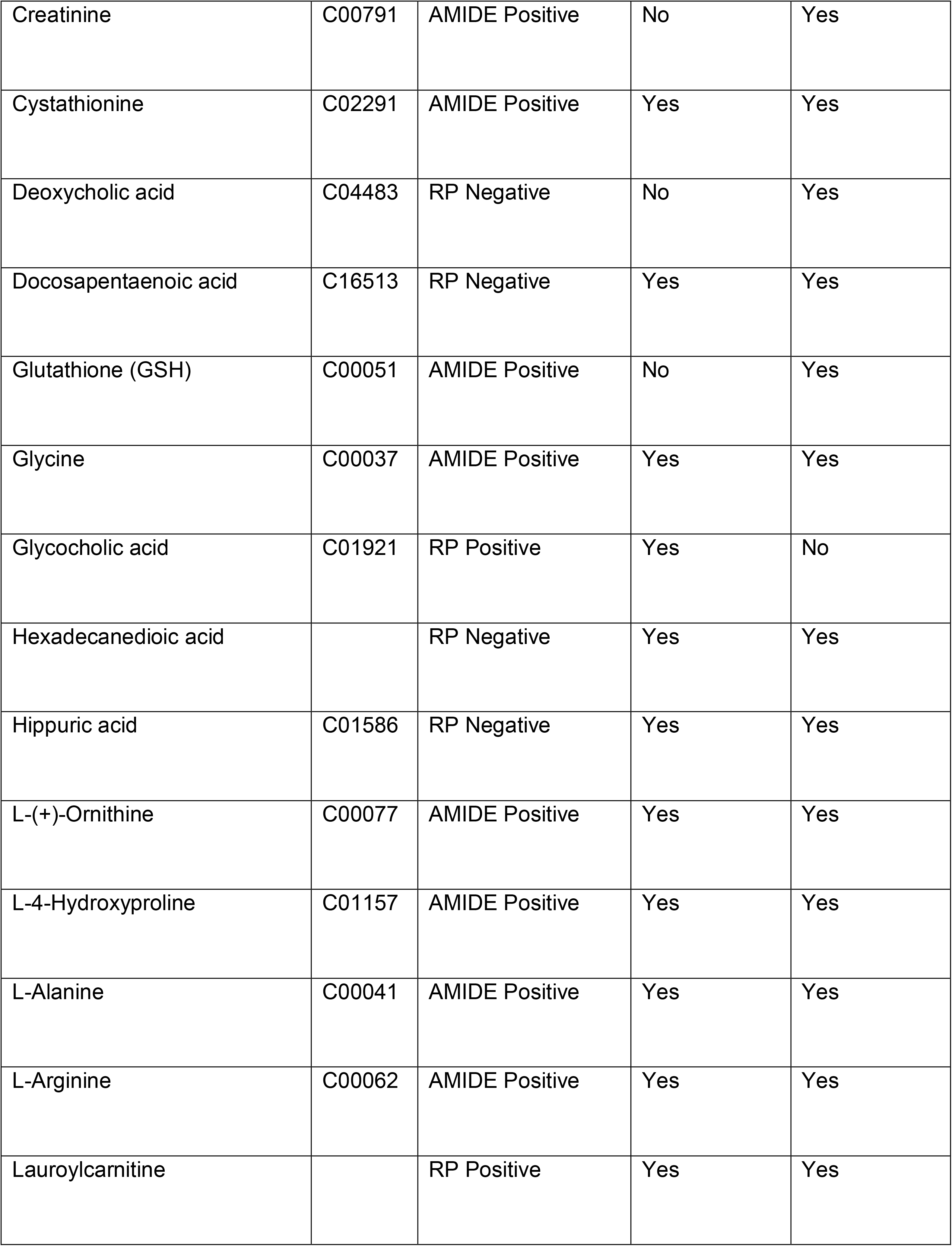
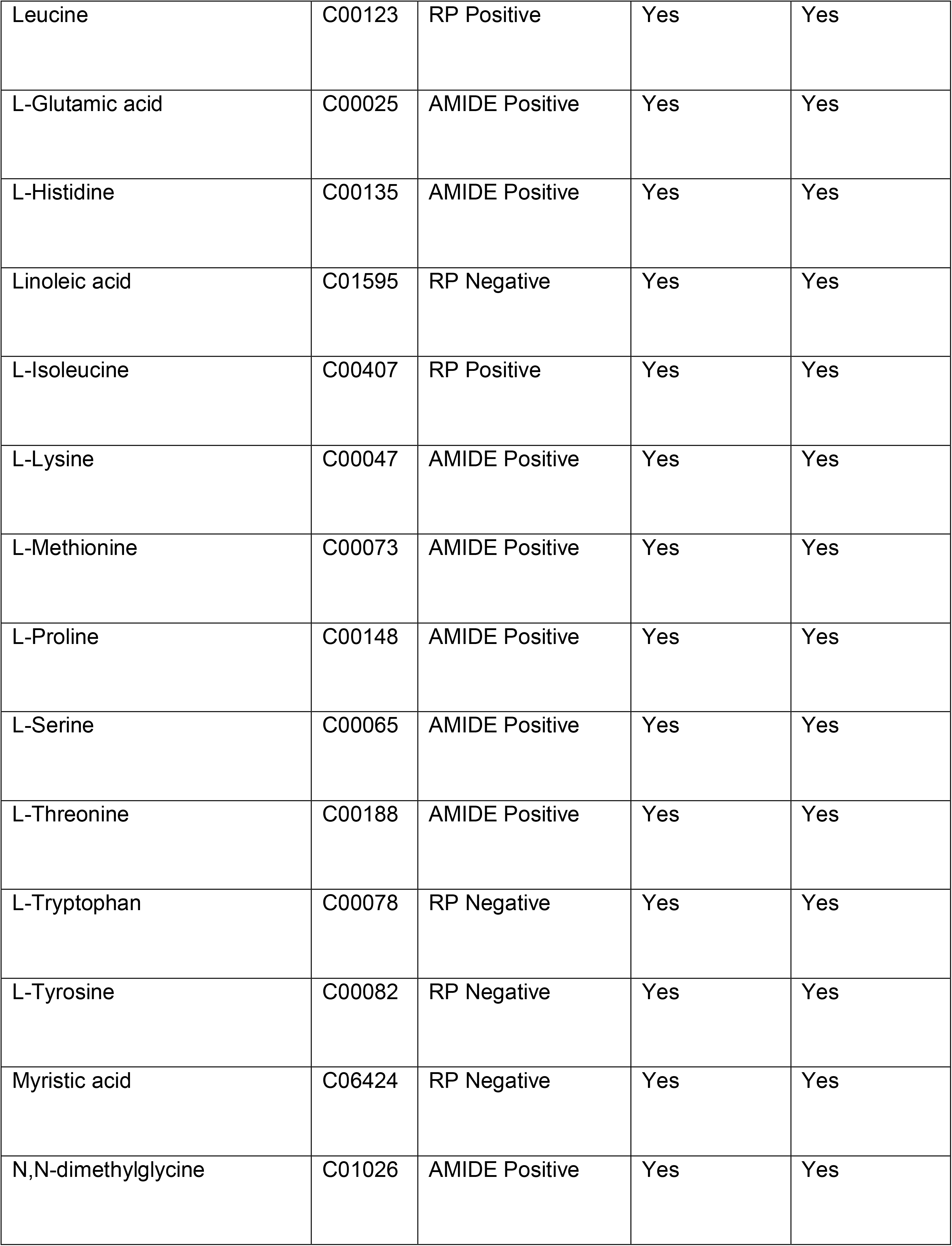
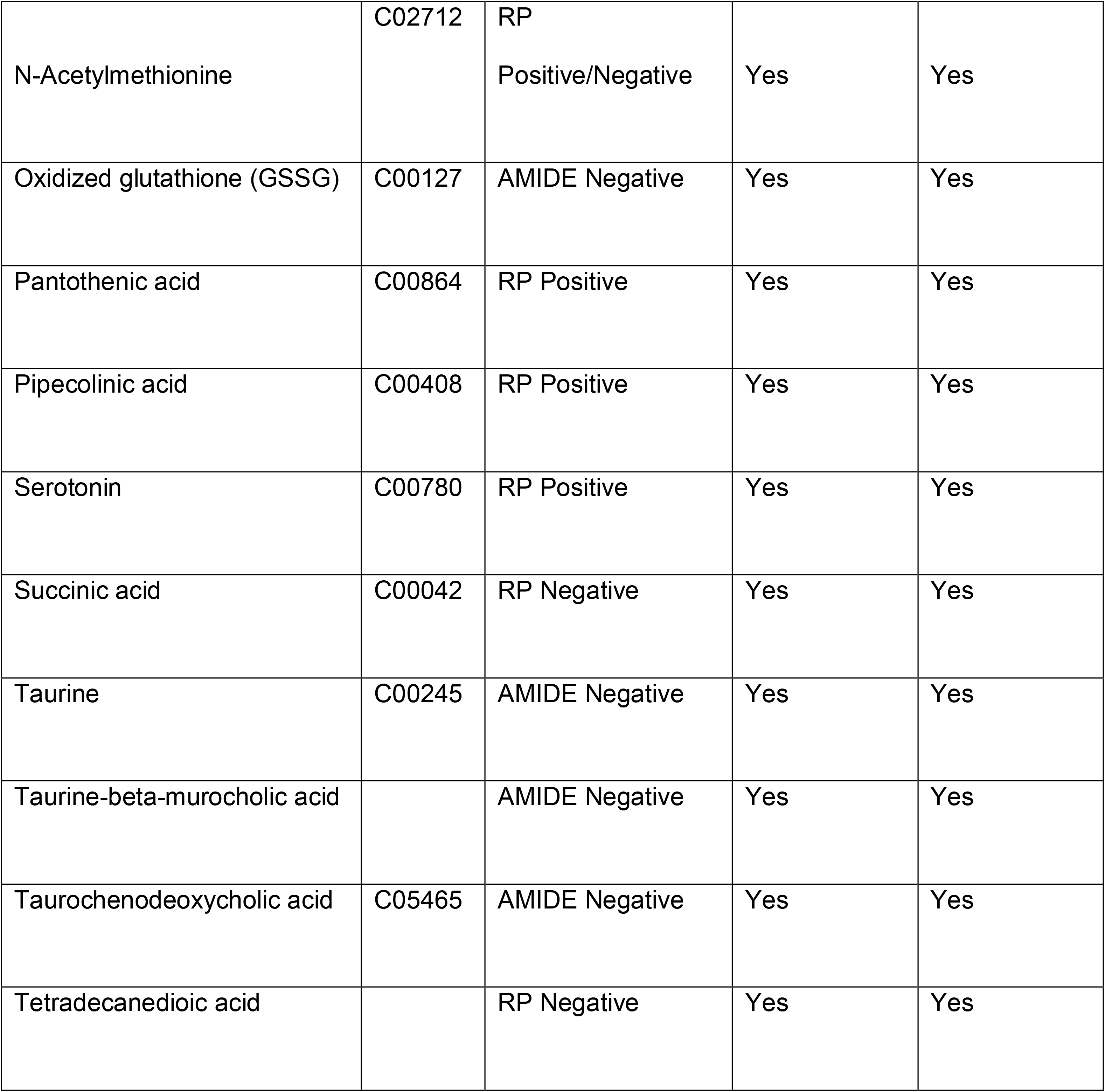
List of metabolites identified by targeted peak extraction in the UPLC/MS data. Table indicates compound name, KEGG Entry number, type of column was used for UPLC and if the peak matched the retention time and MS2 spectra identified with the chemical standard in liver and plasma samples.

